# Selecting 16S rRNA primers for microbiome analysis in a host-microbe system: the case of the jellyfish *Rhopilema nomadica*

**DOI:** 10.1101/2022.10.17.510620

**Authors:** Noga Barak, Eduard Fadeev, Vera Brekhman, Dikla Aharonovich, Tamar Lotan, Daniel Sher

## Abstract

Amplicon sequencing of the 16S rRNA gene is extensively used to characterize bacterial communities, including those living in association with eukaryotic hosts. Deciding which region of the 16S rRNA gene to analyze and selecting the appropriate PCR primers remained a major decision when initiating any new microbiome study. Here, we compared three commonly used primers targeting different hypervariable regions of the 16S rRNA gene, V1V2, V3V4, and V4V5, using the jellyfish *Rhopilema nomadica* as a model. Although all primers exhibit a similar pattern in bacterial community composition, the performance of the V3V4 primer set was superior to V1V2 and V4V5. The V1V2 primers misclassified bacteria from the Bacilli class and exhibited low classification resolution for Rickettsiales, which represent the second most abundant 16S rRNA gene sequence in all the primers. The V4V5 primer set detected almost the same community composition as the V3V4, but the ability of these primers to also amplify eukaryotic 18S rRNA gene may hinder bacterial community observations. However, after overcoming the challenges possessed by each one of those primers, we found that all three of them show very similar bacterial community dynamics and composition. Nevertheless, based on our results, we propose that the V3V4 primer set is the most suitable for studying jellyfish-associated bacterial communities. Our results suggest that, at least for jellyfish samples, it may be feasible to directly compare microbial community estimates from different studies, each using different primers but otherwise similar experimental protocols. More generally, we recommend specifically testing different primers as a prelude to large-scale 16S rRNA gene amplicon analyses, especially of previously unstudied host-microbe associations.

## Introduction

The existence of prokaryotic microbes dates long before the first eukaryotic animals appeared and it is likely that host-bacteria symbioses have existed for hundreds of millions of years (Bosch, 2013; Weiland-Bräuer et al., 2020). These symbioses are not only important and can influence health and disease of many hosts, but they can also help us understand evolution and development process (Fraune and Bosch, 2010; Foster et al., 2017). In recent year, alongside studies of bacteria as pathogens, there is an increasing evidence on bacteria playing significant roles in shaping the phenotype, development, behavior, and fitness of various hosts (Gould et al., 2018; Rosenberg and Zilber-Rosenberg, 2018; Schnedler-Meyer et al., 2018; Baldassarre et al., 2021). The host associated microbial community can provide numerous beneficial functions, including nutrition, assisting in the maturation of the immune system or protection from pathogenic infections (Mortzfeld et al., 2018; Stabili et al., 2018a). These important roles played by symbiotic bacteria in the lives of extant eukaryotic hosts raise the question of how these symbioses may have contributed to host evolution, as postulated by the hologenome theory of evolution (Zilber-Rosenberg and Rosenberg, 2008; Rosenberg and Zilber-Rosenberg, 2011). Answering such questions could be facilitated by the study of ancient phyla such as cnidarians and their associated symbionts.

Cnidarians, which include sea anemones, corals, hydrozoans, jellyfish and parasitic myxozoas, are one of the oldest animal phyla, having evolved 700 million years ago (Cartwright et al., 2007). Microbes associated with corals, hydra, jellyfish and other cnidarians have been extensively investigated since they impact the health and fitness of the cnidarian host (Fraune and Bosch, 2010; van Oppen and Blackall, 2019). These studies suggest that the cnidarian microbiome differs between closely related organisms, is distinct from that of the surrounding water, and varies as the life cycle of the host progresses, e.g. from benthic to pelagic (Tinta et al., 2019). Several studies have also shown differences in microbial community structure between healthy and diseased cnidarian tissue, thus linking microbiome structure with the health of the host (Stabili et al., 2018b; Van De Water et al., 2018). Given the importance of cnidarians (especially corals and some hydrozoa) as reef-building organisms, and the prevalence of jellyfish blooms worldwide, understanding how the cnidarian microbiome is related to host health is important, e.g. for conservation or mitigation efforts (Peixoto et al., 2017; Tinta et al., 2019; Voolstra et al., 2021). However, research on structure of cnidarian-associated bacteria is challenging (Apprill 2017) and is currently conducted using highly diverse methodologies. The technical discrepancies between the studies impede the establishment of general overview on the cnidarian microbiome.

The most commonly used methodology for taxonomic profiling of bacterial communities is amplification and sequencing of (part of) the 16S rRNA gene. Since the 16S rRNA gene consists of nine variable regions which can be targeted by different primers (in different combinations) for sequencing, choosing the right primer combination is important (Figure 1A). For instance, The Earth Microbiome Project, which aims to characterize and interpret the microbial diversity and functional potential of thousands environmental samples, uses primers that target the V4 hypervariable region (Thompson et al., 2017). In contrast, the Human Microbiome Project uses mostly primers spanning the V1V3 and V3V5 regions (Methé et al., 2012), some studies focusing on clinical studies and the gut microbiome employ primers spanning the V3V4 and V1V2 regions (Alcon-Giner et al., 2017; Liu et al., 2020), and many studies of the ocean target the V4V5 region (Parada et al., 2016). Even within a more limited range of environments or hosts, such as in studies of cnidarian microbiomes, there is not standard primer set used in all studies. Since 2015, 152 studies investigated the taxonomic composition of cnidaria-associated bacteria, using 11 different primer sets, each targeting different regions of the 16S rRNA gene (Figure 1B). Therefore, the question of whether there are “optimal” primer sets for different environments or hosts, including cnidarians, remains open (Klindworth et al., 2013; Yang et al., 2016; Wear et al., 2018; Willis et al., 2019; Fadeev et al., 2021).

**Figure 1.**
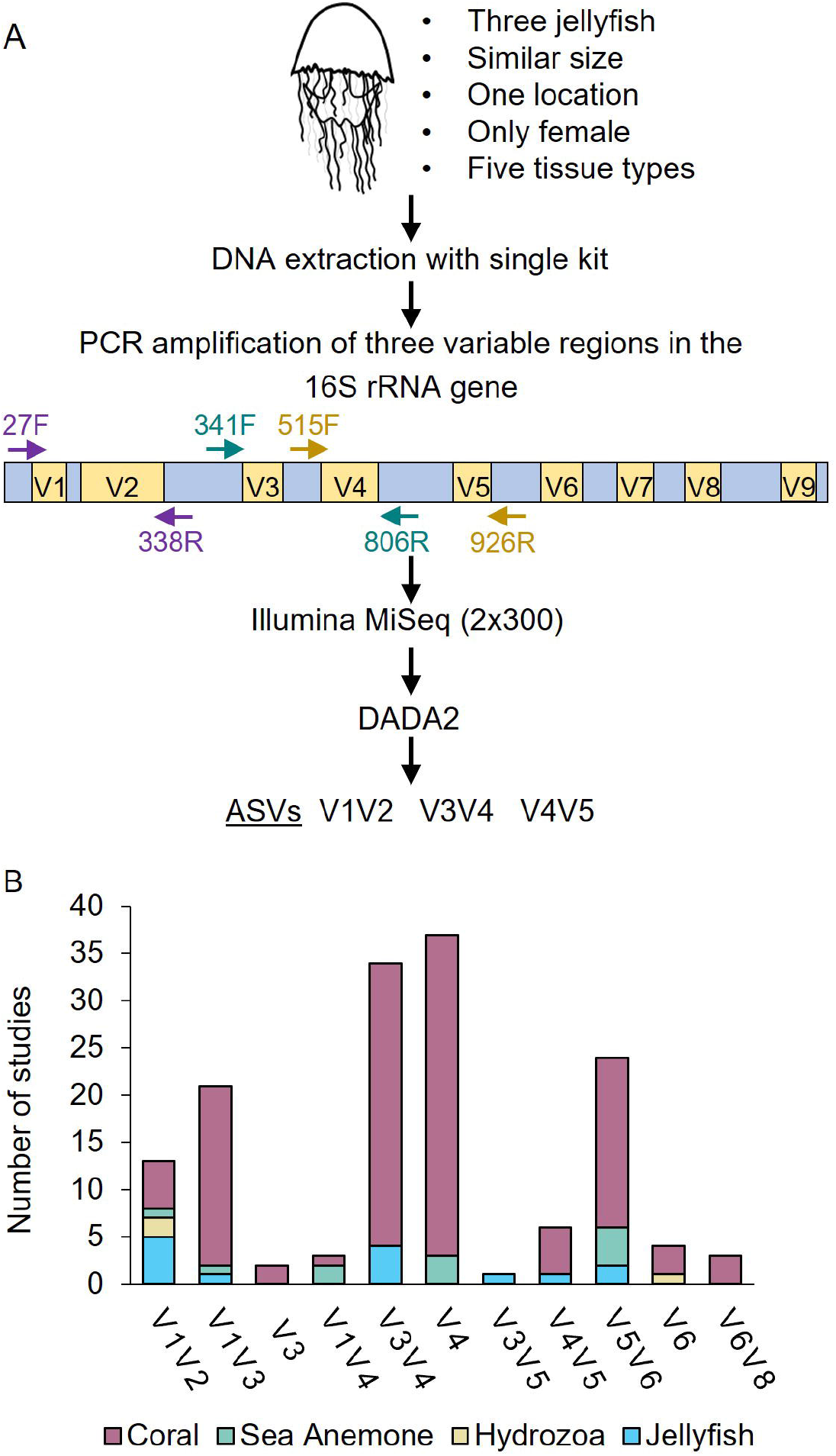
Overview of analysis stages used in this study. A) Schematic illustration of the processing stages. DNA from jellyfish tissues was extracted. Amplicons were generated using different primer pairs targeting different variable regions and sequenced on an Illumina MiSeq. Afterwards, the sequnces went through the same bioinformatic workflow on DADA2 and ASVs tables were created for each primer. B) Number of studies and primer sets types used for Cnidaria microbiome characterization since 2015.

Our model jellyfish is *Rhopilema nomadica*, which forms massive blooms with significant ecological and economic impacts (Lotan et al., 1992; Angel et al., 2014, 2016; Edelist et al., 2020). To select the most suitable 16S rRNA primer set for a large, multi-annual study of *R. nomadica* microbiome in the Eastern Mediterranean, we compared the performance of three commonly used 16S rRNA primer sets (V1V2, V3V4, and V4V5). We examined the performance of the different primer sets in terms of the number of useful sequences, assessments of diversity and the taxonomical assignment as individual ASVs (Amplicon Sequence Variants). Furthermore, we asked whether it is possible to combine the results of the different primer sets. While our analysis was focused on a jellyfish, *R. nomadica*, we believe the systematic comparison between different 16S rRNA primers can provide useful insights for the study of host-associated bacteria in general.

## Materials and methods

### Jellyfish and sea water samples collection and preparation

On the 19th of July 2020, during a major jellyfish bloom in the eastern Mediterranean, three female *Rhopilema nomadica* jellyfish of similar sizes (24-32 cm) were collected from Haifa Bay, Israel (32°50’22.9”N 35°00’04.8”E). The jellyfish were caught individually by swimmers using round buckets to avoid damage to the animals and then gently transferred to 80-liter containers on the boat filled with ambient seawater. All the collected jellyfish were active and healthy. Animal dissection and sample preservation were performed less than two hours after specimen collection. The diameter of each jellyfish was measured, and different tissues were collected from each jellyfish-bell, tentacles, gastrovascular canals, gonads, and rhopalium, using sterile tool kits. After the collection of each tissue, the tools went through second process of cleaning using 1% sodium hypochlorite, DNA AWAY™ (Thermo), 70% Ethanol and Ultrapure (miliQ) Water. Triplicates from each tissue were collected and placed immediately on dry ice. Upon arrival at the lab, the frozen tubes were kept in - 80°C for later use. To identify the gender of the jellyfish, gonad samples from each jellyfish were maintained on ice and observed using a dissecting microscope (Zeiss Axio Imager M2). In addition to the jellyfish samples, five liters of seawater were filtered on a Sterivex filter cartridge (0.22 μm), 1ml of preservation/lysis solution was added (40mM EDTA, 50mM Tris pH 8.3, 0.75M Sucrose) and the samples were frozen as described above.

Bell, gastrovascular canals, gonads, and rhopalium samples were homogenized using a bead beater (TissueLyser II Qiagen) followed by 1h treatment with lysozyme at 37ºC (Merck, 100 mg/ml) and 1h proteinase k at 55 ºC (Promrga, 20 mg/ml). DNA of the total microbial community was extracted with ZymoBIOMICS DNA Miniprep Kit (Zymo Research) following the manufacture protocol. The Sterivex filter (Millipore) was cut as described by Cruaud et al., 2017 and the filter was placed in the ZR BashingBead™ Lysis Tubes from the ZymoBIOMICS DNA Miniprep Kit. Subsequent DNA extraction was performed as described above for the tissue samples.

For standardization, a commercially available mock microbial community standard was used (ZymoBIOMICS™, Zymo Research). The mock community was extracted using 75 µl per prep as recommended by the manufacturer following the same protocol as all other samples.

### 16S rRNA gene library preparation and sequencing

Three different hypervariable regions of the bacterial 16S rRNA gene were amplified using aliquots of the isolated DNA from each sample-V1V2, V3V4, and V4V5 (Supplementary Table S1). Amplicons were generated using a two-stage polymerase chain reaction (PCR) amplification protocol as described previously (Naqib et al., 2018). The primers contained 5’ common sequence tags (known as common sequence 1 and 2, CS1 and CS2). First stage PCR amplification was carried out using DreamTaq Green PCR master mix (M/s Thermo scientific, USA). Briefly, each 50µl reaction mix contained 25µl DreamTaq Green PCR master mix (2X), 1µl (10 µM) each of forward and reverse primers, 2 µl of template DNA and 21 µl of nuclease-free water. The amplification parameters were set as follows: 95° C for 5 min, followed by 32 cycles at 95°C for 30s, 52°C for 45s, and 72°C for 1 min, and a final extension phase of 72°C for 7 min. One microliter of PCR product from the first stage amplification was used as template for the 2^nd^ stage, without cleanup. Cycling conditions were 98°C for 2 minutes, followed by 8 cycles of 98°C for 10s, 60°C for 1min and 68°C for 1min. Libraries were then pooled and sequenced with a 15% phiX spike-in on an Illumina MiSeq sequencer employing V3 chemistry (2×300 base paired-end reads). Library preparation and sequencing were performed at the Genomics and Microbiome Core Facility (GMCF; Rush University, IL, USA).

### Literature survey of microbiome studies in cnidaria

Publications were searched on PubMed using the following query: (16S rRNA) AND (“bacterial composition” OR microbiome OR microbiota) AND (cnidaria OR jellyfish OR Sea Anemone OR hydra OR hydrozoa OR coral OR myxozoa). Only publications between the years 2015 – 2022 were selected. The query resulted in 232 papers where 152 were selected after filtration. The papers we filtered out included bacteria isolation reports or studies on non-cnidaria hosts that were associated with coral reefs. A complete list of papers and details on each specific term are in Supplementary Table S5.

### Data analysis and visualization

Quality control of the raw paired-end reads was performed before the analysis using FastQC v0.11.9 tool. Further analysis was conducted using R 4.1.0 in RStudio v1.4.1717-3. All of the amplicons libraries were processes using DADA2 (Callahan et al., 2016a) and following the recommended workflow (Callahan et al., 2016b). The forward reads were trimmed to 220bp (V1V2), 240bp (V3V4), or 260bp (V4V5) and the reverse reads to 200bp (V3V4) or 215bp (V3V4 and V1V2). Forward and reverse reads were merged based on minimum overlap of 8 bp, chimeras were filtered out and amplicon sequence variants (ASVs) table was created. The representative ASVs were taxonomically classified against SILVA 16S rRNA gene reference database v138.1 (Yilmaz et al., 2014). ASVs that were not classified at the phylum level or were not assigned to bacterial lineages (including those assigned to mitochondria and chloroplasts), were excluded from further analysis.

Over management of the data (organization of tables) was performed using the R package “phyloseq” v1.40.0 (McMurdie and Holmes, 2013). Plots were generated using the R package “ggplot2” v3.3.6 (Wickham, 2016). Alpha diversity indexes were calculated using the function “estimate_richness” in “phyloseq”. Kruskal-Wallis rank sum test was done using R package “rstatix” v0.7.0 (Kassambara, 2021). Non-metric multidimensional scaling plots were created to determine differences in the bacteria communities of different primers sets using R packages “phyloseq” v1.40.0 (McMurdie and Holmes, 2013) and “vegan” v2.6.2 (Callahan et al., 2016b). For the NMDS, Bray-Curtis distance between samples was calculated based for relative abundance values of sequences (ASVs) and genus levels. Two-way permutation multivariate analysis of variance (“Two way PERMANOVA”) was conducted using the R package “vegan” v2.6.2 (Oksanen et al., 2022). Bar plots and Venn diagrams were created using the R packages dplyr v1.0.9 (Müller, 2022) and “VennDiagram” v1.7.3 (Hanbo Chen, 2011), respectively.

Sequences of different molecular mass bands obtained with V4V5 primers were uploaded and processed via SILVAngs v.1.9.8/1.4.9 (https://www.arb-silva.de/ngs/). Visualization of the relative abundances the different molecular mass bands preformed using Krona RSF display tool (Ondov et al., 2011). In some cases, relevant ASV sequences (table S3) were extracted and classified using SINA against SILVA database and BLAST against the NCBI databases (Johnson et al., 2008; Quast et al., 2013).

## Results

### Multiple amplicon sequence lengths affect community coverage

PCR amplification of the 16S rRNA gene was expected to result in a single band (∼355bp, ∼510bp, ∼450bp for the V1V2, V3V4 and V4V5 primer sets, respectively). Indeed, the PCR results of V1V2 and V3V4 produced single bands of the expected size (∼355bp, ∼510bp respectively). In contrast, a double band was observed in most of the samples using the V4V5 primers (∼450bp and ∼600bp, Figure 2A). The V4V5 primer set has previously been shown to amplify also the 18S gene of eukaryotes (Parada et al., 2016; Parada and Fuhrman, 2017; Yeh et al., 2021, see discussion below), and indeed sequencing of DNA from each band extracted from the gel separately shows that the high molecular mass band corresponded to amplified jellyfish 18S rRNA gene, and the low molecular mass amplicon corresponded to amplified 16S rRNA from the prokaryotic microbial community (Fig 2B). The ratio between the intensities of the two PCR bands in the V4V5 amplicons varied between samples and jellyfish tissues without any clear pattern. Due to the length of the V4V5 18S eukaryotic amplicon, the merging procedure for paired reads in DADA2 was largely unsuccessful, resulting in far fewer usable sequences for this primer set compared to the others (Fig 2C).

**Figure 2.**
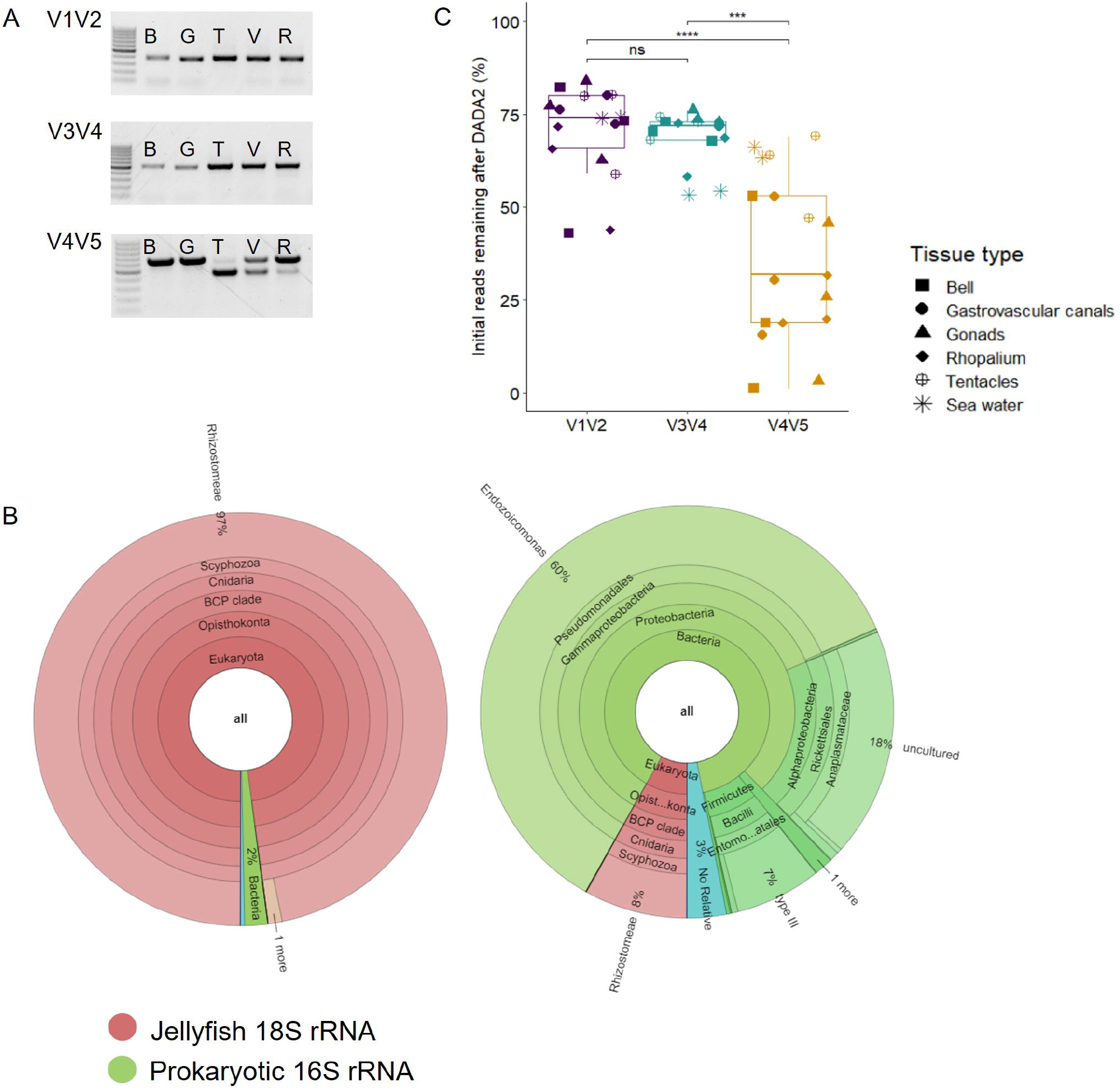
Double bands effect on downstream analysis. A) PCR amplicons products for five jellyfish tissues, and three primers set. B- bell, G- gonads, T- tentacles, V- gastrovascular canals, R- rhoplium. B) Percentage of initial reads remaining after DADA2 processing.V4V5 remining reads were significant less than the other two primer sets (Kruskal-Wallis and Dunn’s tests. p<0.001) C) Visualization of relative abundances sequences in high vs low bands obtained with V4V5 primers with Krona RSF display tool (Ondov et al., 2011).

In addition to the two clearly discernable amplicons in the V4V5 primers, there was also heterogeneity in the amplicon length (after trimming and merging paired ends) which differed between the primer sets. We initially noticed this heterogeneity in the mock community, where using a window of 300-320 bp with the V1V2 primers resulted in identification of only 7 out of the 8 known community members. Subsequent analysis of the range of sizes in the various primer sets revealed differences in their size range (273-343 bp for V1V2, 400-430 for V3V4 and 366-378 for V4V5). These differences were biologically relevant, as the smallest amplicons in the V1V2 primers were from *Synechococcus* (abundant seawater cyanobacteria), and the largest from *Lactobacillacea* from the Mock community. The sequences corresponding to these clades would have been missed if stringent size criteria were used to select the sequences for further analysis.

### Similar pattern of diversity and community structure revealed by all three primer sets

All the primer sets revealed similar patterns in diversity. The total ASV number and the Simpson index showed no significant differences between the primer sets (Figure 3, ANOVA, p>0.05). In all primers, the highest diversity was observed in the seawater and the bell tissues compared to the other tissues. In each of the primer sets, the seawater samples clustered separately from the jellyfish samples (Figure 4A) and within the jellyfish samples, the bells clustered apart from all other tissues. The grouping of the samples corresponded to the tissue type, rather to the individual jellyfish (Figure 4A, PERMANOVA test; differences between tissue type p-value <0.001 for all primers, differences between individual jellyfish p-value >0.2).

**Figure 3.**
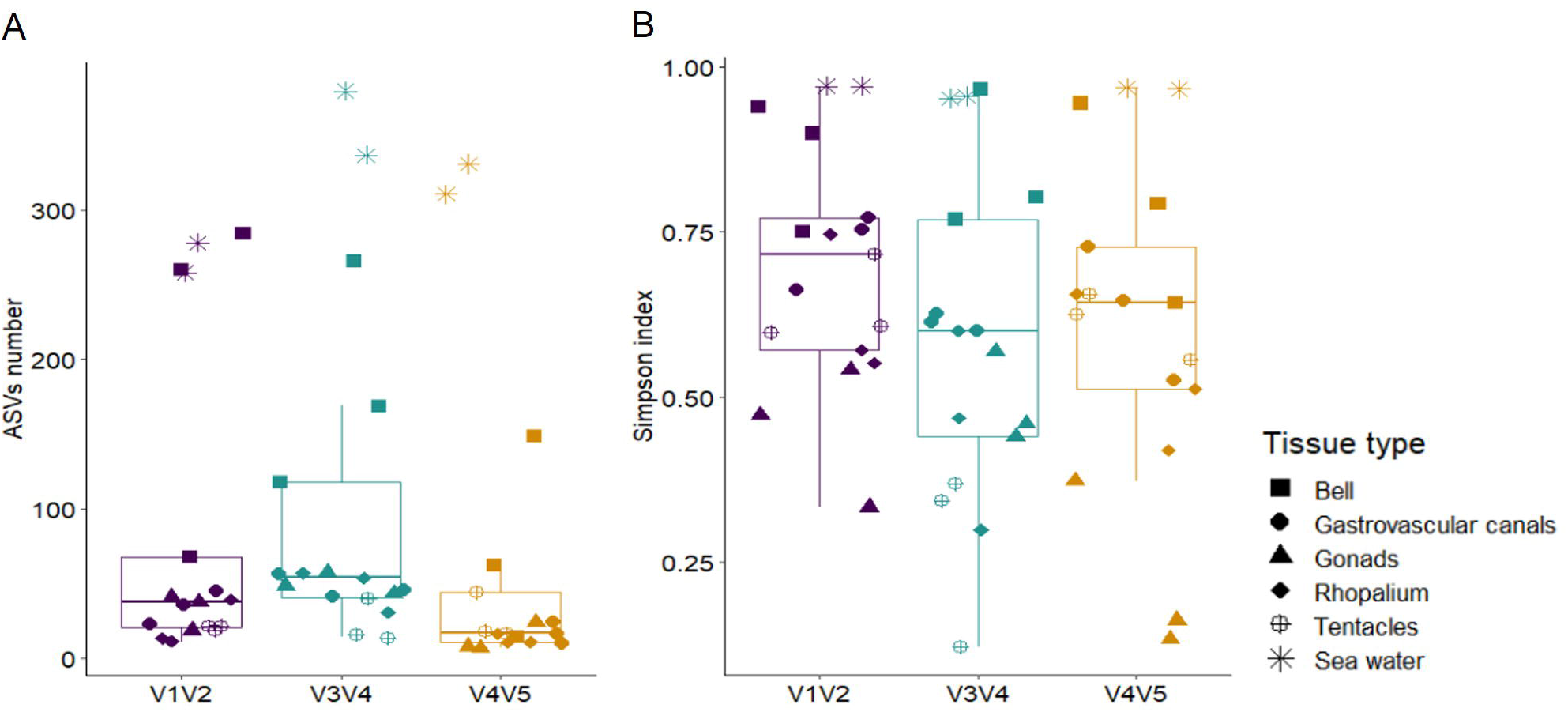
Alpha diversity indexes. A) ASVs numbers. B) Simpson’s Diversity Index. On both alpha diversity indexes no significant differences were seen between primers (ANOVA, p>0.05)

**Figure 4.**
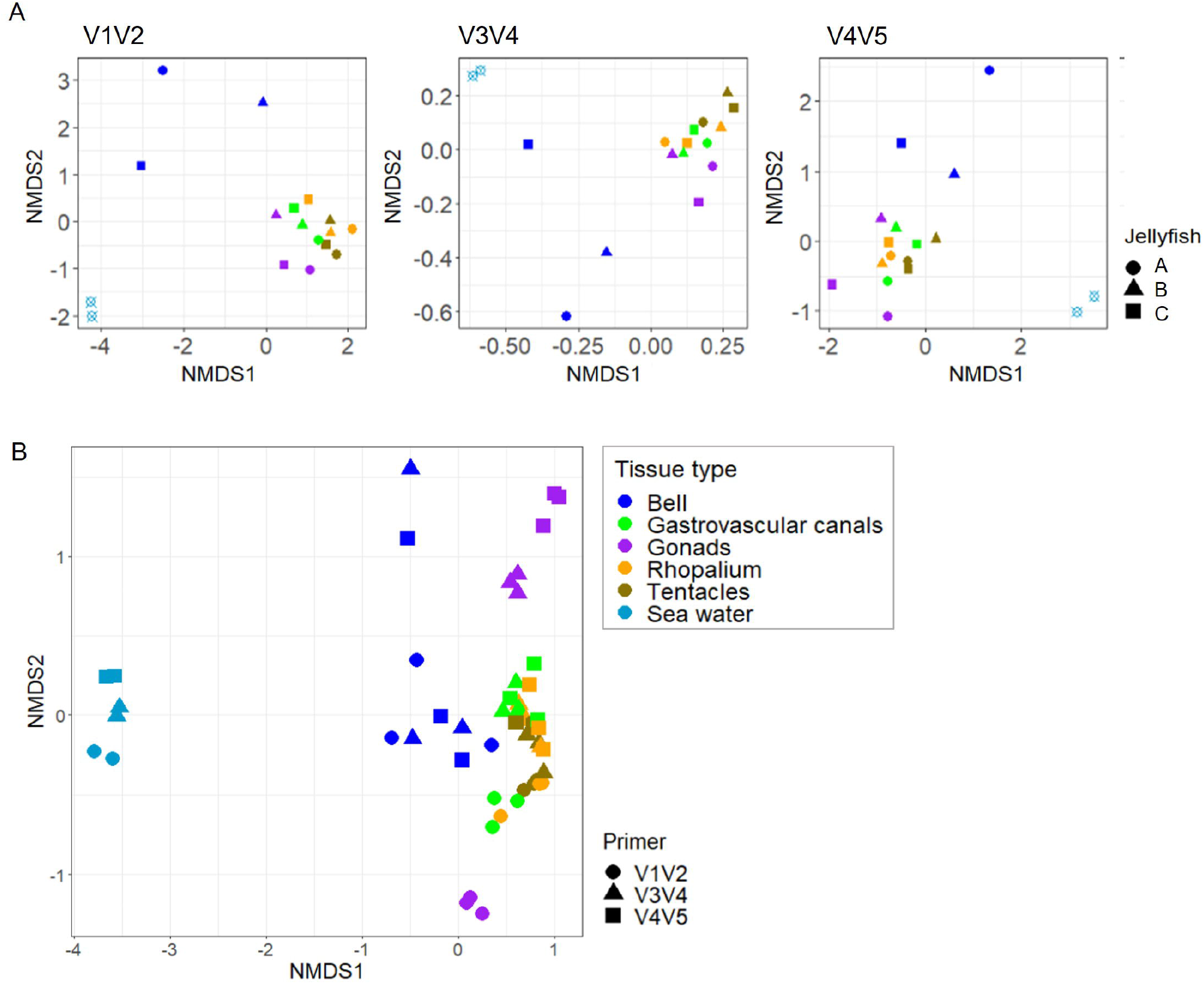
Non-metric multidimensional scaling (NMDS) of the bacterial community composition in different jellyfish and tissues obtain with different primers. A) ASVs based NMDS of V1V2 primer (stress 0.07), V3V4 primer (stress 0.08) and V4V5 primer (stress 0.09). B). Taxonomy based NMDS analysis. Bacterial community composition of both ASVs and taxonomy based NMDS are based on Bray-Curtis dissimilarity (stress 0.10).

As the regions of the 16S rRNA gene amplified by each primer set do not overlap, the resulting ASVs cannot be compared directly (e.g., aligned together). Therefore, to determine whether there are major differences between the primer sets in the observed microbial community, following Fadeev et al., (2021), we compared the datasets from the three primers based on the sequence counts at the genus rank (i.e., the lowest shared taxonomic rank). Sequences that were unclassified at the genus rank, were defined at the lowest classified rank (i.e., phylum, class, order, or family). Overall, the ASVs were merged into 454 different genera and another 147 lineages that could not be classified at the genus level. The three primer sets shared 151 of the total 454 genera, with 50-143 genera unique to each primer set (Figure 5A). The shared 151 genera represent the vast majority of the 16S sequences in each dataset (>94%), and thus, the observed microbial communities (Figure 5B).

**Figure 5.**
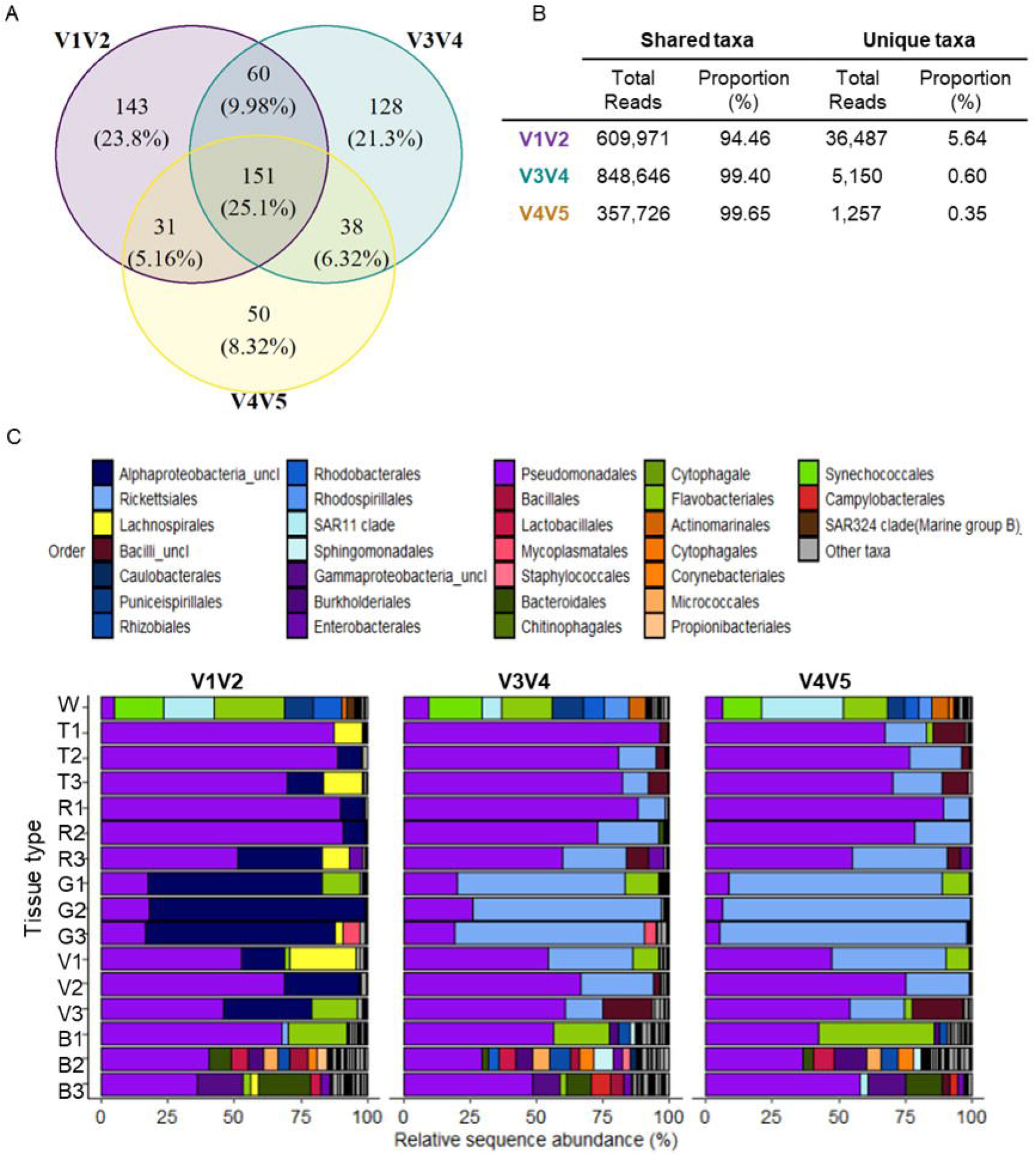
V1V2, V3V4 and V4V5 merged datasets analysis. A) Venn diagram representing all genus from the merged datasets. B) Reads numbers and proportion of shared and unique genera compared to the complete population of each primer. C) Taxonomic compositions (relative sequence abundance) of order level in different jellyfish tissues (B- bell, G- gonads, T- tentacles, V- gastrovascular canals, R- rhoplium) and one sample of filtered seawater (W). Taxa with relative sequence abundance of less than 2% were classified as “Other taxa”. Taxa discussed in the text are at the top of the bars.

The dissimilarly patterns of the combined dataset remained similar to the ASV-based datasets of individual primer sets (Figure 4B). This differences between the tissue types were observed primarily along NMDS1, whereas a clear portioning could be seen along NMDS2 between the V1V2 primers (excluding the bell tissue) and the two other primer sets (“mirror image”, Figure 4B). The PERMANOVA test showed there is a significant difference (p-value < 0.001) of both primers and tissues type. However, differences between tissues were correlated with 64% of the variability whereas the different primers represented less than 10% (R2= 0.64 and 0.099 respectively). Therefore, while there are differences between the primer sets, to a large extent these do not mask the biological variability between the samples. (PERMANOVA test; R 2 = 0.099, p-value < 0.001; Figure 4B). Therefore, while there are differences between the primer sets, to a large extent these do not mask the biological variability between the samples.

### Discrepancies in taxonomic classifications between the primers sets

We next determined whether there were specific lineages identified differently by each primer set, which could explain the different ordination of the V1V2 primer amplicons in Figure 4B. One of the most abundant shared lineages (21-23% of the sequences) could be classified by the V3V4 and V4V5 primers to the order level, whereas it could only be classified to the phylum level by the V1V2 primer set (Rickettsiales vs. Alphaproteobacteria, dark and light blue, respectively, in Figure 5C).

In addition to differences in the taxonomic resolution of ASV classification, we also encountered a case where the same organism was classified by the different primers as belonging to two different classes, resulting in incorrect identification of a “primer-specific” lineage. Specifically, the fraction of unique sequences was much higher for the V1V2 primer set compared with the two others (ca. 6% compared with less than 1% of all sequences in each dataset, respectively; Figure 5B). Most of the unique sequences in V1V2 dataset (4.5%) corresponded to the 4^th^ most abundant sequence on the dataset and were classified as “unclassified genus” of Lachnospiraceae (yellow in Figure 5C). In both V3V4 and V4V5 primers dataset, the 3^rd^ most abundant sequences represent a similar fraction of the community (4-4.6%) and were classified as “unclassified Bacilli” (brown in Figure 5C). To clarify whether Lachnospiraceae and “unclassified Bacilli” reads originate from the same organisms, we extracted the relevant ASV sequences from all primer sets and classified them against both SILVA and NCBI databases using BLAST. The manual BLAST with the V1V2 and V4V5 ASV sequences resulted in classification to the genus level as Spiroplasma (Firmicutes-Bacilli-Entomoplasmatales-Spiroplasmataceae-Spiroplasma), whereas the V3V4 classification was only to the order level (Entomoplasmatales). This was in agreement with the previous results for V3V4 and V4V5 (as part of the Bacilli class), and in contrast with the V1V2 classification as Lachnospiraceae. We note that the identity level to the best hit in the SILVA database was only 82.7-87.8%. However, BLAST against the NCBI database resulted in a much higher identity level, and all the sequences were identified as Spiroplasma (99.5-100%. Supplementary Table S4).

## Discussion

The sequencing of short PCR amplicons from the 16S rRNA gene (or its RNA product) is an efficient and cost-effective way to characterize the composition of bacterial communities across niches, including environmental and host-associated communities. Despite the high popularity of 16S rRNA amplicon sequencing, choosing the appropriate region of the 16S rRNA gene for sequencing is not trivial and the choice should be done thoughtfully. In some environments, detailed benchmarking studies have compared different primer sets (e.g., Beckers et al., 2016; Graspeuntner et al., 2018; Willis et al., 2019), however such studies are lacking in cnidarians, and are rare in marine host-associated bacteria in general. Here we investigated the performance of three frequently used primer sets, which target the V1V2, V3V4, and V4V5 regions of the 16S rRNA gene, in representing the taxonomic composition of bacterial communities associated with the jellyfish *R. nomadica*. Based on our results, we have decided for the V3V4 primer set, as the most suitable choice for our long-term studies on *R. nomadica*. Below we discuss the identified performance characteristics of each primer set, focusing on *R. nomadica* but extending, when possible, to other organisms or ecosystems. We then ask to what extent data from multiple primers sets is comparable.

### Amplification of eukaryotic 18S rRNA gene by the V4V5 primer set may affect bacterial diversity observations

The V4V5 primer set is extensively used in research on pelagic microbial communities (e.g. Parada et al., 2016; Mestre et al., 2018; Fadeev et al., 2021). This primer set was initially designed to appropriately amplify the 16S rRNA gene of the highly abundant SAR11 clade (Apprill et al., 2015; Parada et al., 2016). Further improvements of the primer set allowed its use also for investigation of eukaryotic microbial communities, through amplification of the 18 rRNA gene (Parada et al. 2016). We therefore tested them in this study, hoping they would facilitate future comparisons between the jellyfish and oligotrophic seawater samples, for example to identify to what extent bacteria from the surrounding seawater colonize the jellyfish. However, samples amplified with the V4V5 primer set produced a “double band” – two differently sized amplicons corresponding to bacterial 16S and eukaryotic 18S rRNA genes. The ability of V4V5 primer to simultaneously amplify bacteria, eukaryote and archaea has been described before, and has led to their use as “three-domain primers” (Needham and Fuhrman, 2016; Parada et al., 2016; Parada and Fuhrman, 2017; McNichol et al., 2021). Recently, Yeh and co-authors (Yeh et al., 2021) developed bacterial 16S and eukaryotic 18S rRNA mock communities and proposed a workflow designed for concomitant analysis of both in which the 18S amplicon data are analyzed without merging the paired ends. However, in our case the strong but non-systematic amplification of eukaryotic jellyfish host DNA caused a major loss of data during initial stages of analysis (i.e., far fewer assembled bacterial sequences Fig 2B, up to an 99% loss). We also speculate that such uneven amplification of host DNA may introduce biases between samples, although our dataset was not large enough to test this possibility. Therefore, based on our results, we chose not to use these primers for our large-scale jellyfish study. We recommend that, if such primers are considered for use in analyzing host-associated microbiomes (e.g., to study also micro-eukaryotic components of the microbiome), their specificity and efficiency should be tested in small scale prior to full-scale analysis.

### Sequences of the V1V2 region may lead to an inaccurate taxonomic classification

We chose to test the V1V2 primers in our study as they are often used in analyses of the human gut microbiome (e.g. (Miyake et al., 2015; Thompson et al., 2015; Allali et al., 2017; Zhang et al., 2017), and have also been used in microbiome studies of different Cnidaria, such as corals, hydrozoa, sea anemones and jellyfish (Figure 1B). In our analysis we observed three technical issues that may have biological implications. Firstly, the size distribution of the V1V2 PCR amplicons was larger than that of the other primer sets. Amplicon length variation of V1V2 and bimodal size distribution for the V3V4 16S regions were reported in the past (and was seen in our data as well), however the data is limited (Vargas-Albores et al., 2017). The large size variation should be considered when using these primers, to make sure that the bioinformatics pipeline does not remove biologically relevant community members. Secondly, the V1V2 primers classified the Rickettsiales to a lower phylogenetic resolution in comparison to the other sets of primers. The Rickettsiales are a group of obligate intracellular bacteria that can be parasitic, symbiotic, or commensal with diverse host range (Salje, 2021; Schön et al., 2022). Rickettsiales are known to be pathogenic to many animals including human and can also be found in cnidarians, including corals (G*orgonia ventalina, Orbicella annularis*, and *Orbicella faveolate*), Hydrozoa (*Hydra oligactis*), and other jellyfish (*Cyanea capillata*) (Daley et al., 2016; Klinges et al., 2019). The Rickettsiales were detected in a high relative sequence abundance in most of the tissues and especially in the gonads (where they were ∼60-80% of sequences) and therefore could potentially be an important member of the jellyfish microbiome. While we did not explore this observation systematically, the lower phylogenetic resolution for this clade may limit the usefulness of this primer set in samples where Rickettsiales are abundant.

Based on manual curation, using SILVA and NCBI databases, we found that abundant sequences that were taxonomically classified as Lachnospiraceae (class Clostridia) are likely affiliated to Spiroplasma (class Mollicutes). This was the main reason the V1V2 amplicon sequences differed from the other two primer sets in the NMDS ordination, as manually replacing the genus names resulted in all three primer sets clustering together (Supplementary Figure S2). Lachnospiraceae are considered to be part of the core microbiome of the human gut (Gosalbes et al., 2011; Vacca et al., 2020) all are anaerobic, and some taxa may be involved in intestinal diseases (Vacca et al., 2020). Their identification in jellyfish tentacles, which are likely to be an oxygen abundance niche, was therefore surprising. In contrast, the Spiroplasmataceae family (identified using BLAST from the ostensibly Lachnospiracea ASV sequences) has previously been found in other jellyfish such as *Rhizostoma pulmo* and *Cotylorhiza tuberculate* (Viver et al., 2017; Stabili et al., 2019). The biology of the Spiroplasma marine lineages is still poorly understood, although they were suggested in sea cucumbers and in some terrestrial invertebrates to have diverse defensive capabilities (e.g., production of toxins and immune system recruitment), (He et al., 2018; Ballinger and Perlman, 2019). The reason for the potential misclassification is unclear, but we note that the ASV sequences shared only ∼82-89% identity with the Spiroplasma sequences in the SILVA database. Thus, the actual identity of the “Spiroplasma-like” organism is still unclear. More broadly, potential misclassification will be hard to identify using a single primer set - we looked more deeply into this clade only because of the observed differences between the three primer sets. We therefore recommend including a comparative analysis using several different primers on a small, representative subset of any large-scale amplicon-based microbiome study of a new model system.

### V3V4 primers are commonly used but not without caveats

We included the V3V4 primers (Klindworth et al., 2013) in our comparison as they were adopted in the official Illumina protocol (Illumina, 2013), and had been extensively used in a variety of niches from environmental studies to the human gut microbiome (Dyksma et al., 2016; Fan et al., 2019; Takagi et al., 2019), as well as in studies of corals and jellyfish (Figure 1B, Supplementary Table S5). Given the challenges described above with the other two primer sets, we chose to use these primers for our ongoing, large scale, analysis of jellyfish-associated bacterial communities. The V3V4 primers are not without their own caveats, however, chiefly their lower amplification efficiency for SAR11, the most abundant heterotrophic clade in large regions of the ocean (Apprill et al., 2015; Parada et al., 2016). In the case of our jellyfish samples, the V4V5 primers did not identify SAR11 as a dominant member of the jellyfish microbiome, and thus we do not expect using these primers to miss an important member of the microbial community. Another drawback of the V3V4 primers is their inability to amplify archaea and micro-eukaryotes, which were not very common in the results of our V4V5 primers from the same jellyfish tissue.

We note that there are several other primer pairs commonly used for cnidarian microbiome analyses, including the V1V3, V4 and V5V6 primer sets (Figure 1). The V1V3 has been used in the human microbiome project (Methé et al., 2012), but we did not include it in our analysis as the V1V2 primers are used more in jellyfish studies. The V4 region is shorter compared to primers that span more than one region and can be sequence using highly cost-efficient platform (e.g., using 2×150 base paired-end reads). It has been endorsed by the Earth Microbiome Project which is large scale survey of environmental microbial communities (Thompson et al., 2017). However several studies have suggested that longer amplicons are preferable (Derakhshani et al., 2016; García-López et al., 2020; Katiraei et al., 2022). The V5V6 primers are considered a good fit for coral microbiome studies (Bayer et al., 2013) but have not been frequently used in other cnidarian systems. Future analyses also including these primers in the comparison may help define an “optimal” set for primers for host-associated microbes.

### Can the results of different primer sets be compared?

Over the last two decades, there have been thousands of studies of microbial ecosystems performed using 16S rRNA amplicons, and much of these data are publicly available. For example, there are ∼150 paper where cnidarian microbiomes have been studied using 16S rRNA gene amplicon sequencing, published since 2015. It would potentially be highly fruitful to directly compare these studies, both in order to identify large-scale trends in microbial populations in space and time, and in order to allow direct comparison between related studies (e.g., different jellyfish species). However, as demonstrated in Figure 1B, these studies often utilize different PCR primers. At least for the specific biological question addressed here (jellyfish microbiomes across tissues), the community composition dynamics did not differ between the three primer sets. Notably, when the taxonomic classification of specific ASVs differed between primer sets, using multiple primer sets allowed for the identification and, potentially, correction of these discrepancies. Taken together, our results suggest that the overall trends observed by the three primers could be directly compared, and merging into a unified dataset is also possible, albeit at a lower phylogenetic resolution (genus rather than ASV). However, it is important to note, that our observations from samples that were processed identically and in parallel (i.e. sample collection, DNA extraction and sequencing), may not be fully relevant for inter-study comparisons, where many other stages of the analysis workflow may introduce biases (reviewed in Pollock et al., 2018; Abellan-Schneyder et al., 2021).

## Conclusions

There are many studies discussing the advantages and disadvantages of different primers, with no consensus (Allali et al., 2017; Pollock et al., 2018; García-López et al., 2020; Abellan-Schneyder et al., 2021; Darwish et al., 2021). Therefore, it is likely that future studies will continue to employ multiple different primers, depending on the specific model system, the preferences of the research team and the importance of comparability with other studies. In studies of *R. nomadica* (and potentially other marine host-bacteria systems), we recommend using the V3V4 primer set. However. because it is difficult to identify any specific primer set that is universally better than others, we recommend performing two steps before initiating any large-scale study of a new model. Firstly, we recommend performing a preliminary literature review of the primers used in related studies or organisms. Secondly, we advise testing several primers on a limited number of samples, to identify (and potentially correct) technical issues, including misclassifications of major members of the microbial community. These stages can also help identify cases where multiple datasets using different primers can be compared.

## Supplementary Data

**Figure S1. Mock community analysis.** Number of ASVs belonging to the mock community vs ASVs sequences lengths. Primers are represented by color.

**Figure S2. Analysis of manual changes of misclassified ASVs in V1V2.** Two ASVs were changed-1. ASV0002 (originally identified as unclassified Alphaproteobacteria 2. ASV0003 (originally identified as Lachnospiracea). A) Taxonomy based NMDS analysis. Bacterial community composition based on Bray-Curtis dissimilarity (stress 0.089). B) Taxonomic compositions (relative abundance) of order level in different jellyfish tissues (B- bell, G- gonads, T- tentacles, V- gastrovascular canals, R- rhoplium) and one sample of filtered seawater (w). Different taxonomic order are represented by color code. Taxa with relative abundance of less than 2% were classified as “Other taxa”.

**Table S1.** Primers used for 16S rRNA gene sequencing

**Table S2.** In silico evaluation of used primers. Coverages were evaluated using Silva TestPrime.

**Table S3.** ASVs used for BLAST analysis. ASV0004 originally assigned taxonomy of unclassified Lachnospiraceae and ASV0003 was originally assigned taxonomy of unclassified Bacilli.

**Table S4.** Manual BLAST results for ASV0003 (V3V4 and V4V5) and ASV0004 (V1V2)

**Table S5.** Literature survey of microbiome studies in cnidaria

## Author Contributions

NB, DS and TL designed the study. NB, DS and TL provided jellyfish and environmental samples for the study. NB conducted the DNA extraction of the samples and the first PCR stage with the help of VB and DA. NB analyzed the data with the help of EF. NB, TL, and DS wrote the manuscript. All authors contributed to the final version of the manuscript.

## Acknowledgments

We thank the many students from the Leon H. Charney School of Marine Sciences, University of Haifa, and others, who helped to collect the jellyfish. We would like to thank Stefan Green and the Rush University Genomics and Microbiome Core Facility (GMCF) team for the sequencing of the samples

## Data Availability Statement

The datasets generated for this study can be found in NCBI PRJNA870285.

## References

Abellan-Schneyder, I., Matchado, M. S., Reitmeier, S., Sommer, A., Sewald, Z., Baumbach, J., et al. (2021). Primer, Pipelines, Parameters: Issues in 16S rRNA Gene Sequencing. mSphere 6. doi: 10.1128/msphere.01202-20.

Alcon-Giner, C., Caim, S., Mitra, S., Ketskemety, J., Wegmann, U., Wain, J., et al. (2017). Optimisation of 16S rRNA gut microbiota profiling of extremely low birth weight infants. BMC Genomics 18, 1–15. doi: 10.1186/s12864-017-4229-x.

Allali, I., Arnold, J. W., Roach, J., Cadenas, M. B., Butz, N., Hassan, H. M., et al. (2017). A comparison of sequencing platforms and bioinformatics pipelines for compositional analysis of the gut microbiome. BMC Microbiol. 17, 1–16. doi: 10.1186/s12866-017-1101-8.

Angel, D. L., Edelist, D., and Freeman, S. (2014). Local perspectives on regional challengesL: jellyfish proliferation and fish stock management along the Israeli Mediterranean coast. doi: 10.1007/s10113-014-0613-0.

Angel, D. L., Edelist, D., and Freeman, S. (2016). Local perspectives on regional challenges: jellyfish proliferation and fish stock management along the Israeli Mediterranean coast. Reg. Environ. Chang. 16, 315–323. doi: 10.1007/s10113-014-0613-0.

Apprill, A., Mcnally, S., Parsons, R., and Weber, L. (2015). Minor revision to V4 region SSU rRNA 806R gene primer greatly increases detection of SAR11 bacterioplankton. Aquat. Microb. Ecol. 75, 129–137. doi: 10.3354/ame01753.

Baldassarre, L., Levy, S., Bar-Shalom, R., Steindler, L., Lotan, T., and Fraune, S. (2021). Contribution of Maternal and Paternal Transmission to Bacterial Colonization in Nematostella vectensis. Front. Microbiol. 12, 1–11. doi: 10.3389/fmicb.2021.726795.

Ballinger, M. J., and Perlman, S. J. (2019). The defensive Spiroplasma. Curr. Opin. Insect Sci. 32, 36–41. doi: 10.1016/j.cois.2018.10.004.

Bayer, T., Neave, M. J., Alsheikh-Hussain, A., Aranda, M., Yum, L. K., Mincer, T., et al. (2013). The microbiome of the red sea coral stylophora pistillata is dominated by tissue-associated endozoicomonas bacteria. Appl. Environ. Microbiol. 79, 4759–4762. doi: 10.1128/AEM.00695-13.

Beckers, B., Op De Beeck, M., Thijs, S., Truyens, S., Weyens, N., Boerjan, W., et al. (2016). Performance of 16s rDNA primer pairs in the study of rhizosphere and endosphere bacterial microbiomes in metabarcoding studies. Front. Microbiol. 7. doi: 10.3389/fmicb.2016.00650.

Bosch, T. C. G. (2013). Cnidarian-microbe interactions and the origin of innate immunity in metazoans. Annu. Rev. Microbiol. 67, 499–518. doi: 10.1146/annurev-micro-092412-155626.

Callahan, B. J., McMurdie, P. J., Rosen, M. J., Han, A. W., Johnson, A. J. A., and Holmes, S. P. (2016a). DADA2: High-resolution sample inference from Illumina amplicon data. Nat. Methods 13, 581–583. doi: 10.1038/nmeth.3869.

Callahan, B. J., Sankaran, K., Fukuyama, J. A., McMurdie, P. J., and Holmes, S. P. (2016b). Bioconductor workflow for microbiome data analysis: From raw reads to community analyses [version 1; referees: 3 approved]. F1000Research 5, 1–48. doi: 10.12688/F1000RESEARCH.8986.1.

Cruaud, P., Vigneron, A., Fradette, M. S., Charette, S. J., Rodriguez, M. J., Dorea, C. C., et al. (2017). Open the SterivexTM casing: An easy and effective way to improve DNA extraction yields. Limnol. Oceanogr. Methods 15, 1015–1020. doi: 10.1002/lom3.10221.

Daley, M. C., Urban-Rich, J., and Moisander, P. H. (2016). Bacterial associations with the hydromedusa Nemopsis bachei and scyphomedusa Aurelia aurita from the North Atlantic Ocean. Mar. Biol. Res. 12, 1088–1100. doi: 10.1080/17451000.2016.1228974.

Darwish, N., Shao, J., Schreier, L. L., and Proszkowiec-Weglarz, M. (2021). Choice of 16S ribosomal RNA primers affects the microbiome analysis in chicken ceca. Sci. Rep. 11, 1–15. doi: 10.1038/s41598-021-91387-w.

Derakhshani, H., Tun, H. M., and Khafipour, E. (2016). An extended single-index multiplexed 16S rRNA sequencing for microbial community analysis on MiSeq illumina platforms. J. Basic Microbiol. 56, 321–326. doi: 10.1002/jobm.201500420.

Dyksma, S., Bischof, K., Fuchs, B. M., Hoffmann, K., Meier, D., Meyerdierks, A., et al. (2016). Ubiquitous Gammaproteobacteria dominate dark carbon fixation in coastal sediments. ISME J. 10, 1939–1953. doi: 10.1038/ismej.2015.257.

Edelist, D., Guy-Haim, T., Kuplik, Z., Zuckerman, N., Nemoy, P., and Angel, D. L. (2020). Phenological shift in swarming patterns of Rhopilema nomadica in the Eastern Mediterranean Sea. J. Plankton Res. 42, 211–219. doi: 10.1093/plankt/fbaa008.

Fadeev, E., Cardozo-Mino, M. G., Rapp, J. Z., Bienhold, C., Salter, I., Salman-Carvalho, V., et al. (2021). Comparison of Two 16S rRNA Primers (V3–V4 and V4–V5) for Studies of Arctic Microbial Communities. Front. Microbiol. 12, 1–11. doi: 10.3389/fmicb.2021.637526.

Fan, L., Wang, Z., Chen, M., Qu, Y., Li, J., Zhou, A., et al. (2019). Microbiota comparison of Pacific white shrimp intestine and sediment at freshwater and marine cultured environment. Sci. Total Environ. 657, 1194–1204. doi: 10.1016/j.scitotenv.2018.12.069.

Foster, K. R., Schluter, J., Coyte, K. Z., and Rakoff-Nahoum, S. (2017). The evolution of the host microbiome as an ecosystem on a leash. Nature 548, 43–51. doi: 10.1038/nature23292.

Fraune, S., and Bosch, T. C. G. (2010). Why bacteria matter in animal development and evolution. BioEssays 32, 571–580. doi: 10.1002/bies.200900192.

García-López, R., Cornejo-Granados, F., Lopez-Zavala, A. A., Sánchez-López, F., Cota-Huízar, A., Sotelo-Mundo, R. R., et al. (2020). Doing more with less: A comparison of 16S hypervariable regions in search of defining the shrimp microbiota. Microorganisms 8. doi: 10.3390/microorganisms8010134.

Gosalbes, M. J., Durbán, A., Pignatelli, M., Abellan, J. J., Jiménez-Hernández, N., Pérez-Cobas, A. E., et al. (2011). Metatranscriptomic approach to analyze the functional human gut microbiota. PLoS One 6, 1–9. doi: 10.1371/journal.pone.0017447.

Gould, A. L., Zhang, V., Lamberti, L., Jones, E. W., Obadia, B., Korasidis, N., et al. (2018). Microbiome interactions shape host fitness. Proc. Natl. Acad. Sci. U. S. A. 115, E11951–E11960. doi: 10.1073/pnas.1809349115.

Graspeuntner, S., Loeper, N., Künzel, S., Baines, J. F., and Rupp, J. (2018). Selection of validated hypervariable regions is crucial in 16S-based microbiota studies of the female genital tract. Sci. Rep. 8, 4–10. doi: 10.1038/s41598-018-27757-8.

Hanbo Chen (2011). Venn Diagram for Five Sets. BMC Bioinformatics 57, 297. doi: 10.2307/2689606.

He, L. S., Zhang, P. W., Huang, J. M., Zhu, F. C., Danchin, A., and Wang, Y. (2018). The enigmatic genome of an obligate ancient Spiroplasma symbiont in a hadal holothurian. Appl. Environ. Microbiol. 84. doi: 10.1128/AEM.01965-17.

Illumina (2013). 16S Metagenomic Sequencing Library Preparation. Amplicon, P. C. R., Clean-Up, P. C. R., Index, P. C. R., 1–28. Available at: http://support.illumina.com/content/dam/illumina-support/documents/documentation/chemistry_documentation/16s/16s-metagenomic-library-prep-guide-15044223-b.pdf.

Johnson, M., Zaretskaya, I., Raytselis, Y., Merezhuk, Y., McGinnis, S., and Madden, T. L. (2008). NCBI BLAST: a better web interface. Nucleic Acids Res. 36, 5–9. doi: 10.1093/nar/gkn201.

Kassambara, A. (2021). Manual Cite.

Katiraei, S., Anvar, Y., Hoving, L., Berbée, J. F. P., and Harmelen, V. Van (2022). Evaluation of Full □ Length Versus V4 □ Region 16S rRNA Sequencing for Phylogenetic Analysis of Mouse Intestinal Microbiota After a Dietary Intervention. Curr. Microbiol. doi: 10.1007/s00284-022-02956-9.

Klindworth, A., Pruesse, E., Schweer, T., Peplies, J., Quast, C., Horn, M., et al. (2013). Evaluation of general 16S ribosomal RNA gene PCR primers for classical and next-generation sequencing-based diversity studies. Nucleic Acids Res. 41, 1–11. doi: 10.1093/nar/gks808.

Klinges, J. G., Rosales, S. M., McMinds, R., Shaver, E. C., Shantz, A. A., Peters, E. C., et al. (2019). Phylogenetic, genomic, and biogeographic characterization of a novel and ubiquitous marine invertebrate-associated Rickettsiales parasite, Candidatus Aquarickettsia rohweri, gen. nov., sp. nov. ISME J. 13, 2938–2953. doi: 10.1038/s41396-019-0482-0.

Liu, P.-Y., Wu, W.-K., Chen, C.-C., Panyod, S., Sheen, L.-Y., and Wu, M.-S. (2020). Evaluation of Compatibility of 16S rRNA V3V4 and V4 Amplicon Libraries for Clinical Microbiome Profiling. bioRxiv, 2020.08.18.256818. Available at: https://www.biorxiv.org/content/10.1101/2020.08.18.256818v1 %0A https://www.biorxiv.org/content/10.1101/2020.08.18.256818v1.abstract.

Lotan, A., Ben-Hillel, R., and Loya, Y. (1992). Life cycle of Rhopilema nomadica: a new immigrant scyphomedusan in the Mediterranean.

McMurdie, P. J., and Holmes, S. (2013). Phyloseq: An R Package for Reproducible Interactive Analysis and Graphics of Microbiome Census Data. PLoS One 8. doi: 10.1371/journal.pone.0061217.

McNichol, J., Berube, P. M., Biller, S. J., and Fuhrman, J. A. (2021). Evaluating and Improving Small Subunit rRNA PCR Primer Coverage for Bacteria, Archaea, and Eukaryotes Using Metagenomes from Global Ocean Surveys. mSystems 6. doi: 10.1128/msystems.00565-21.

Mestre, M., Ruiz-González, C., Logares, R., Duarte, C. M., Gasol, J. M., and Sala, M. M. (2018). Sinking particles promote vertical connectivity in the ocean microbiome. Proc. Natl. Acad. Sci. U. S. A. 115, E6799–E6807. doi: 10.1073/pnas.1802470115.

Methé, B. A., Nelson, K. E., Pop, M., Creasy, H. H., Giglio, M. G., Huttenhower, C., et al. (2012). A framework for human microbiome research. Nature 486, 215–221. doi: 10.1038/nature11209.

Miyake, S., Kim, S., Suda, W., Oshima, K., Nakamura, M., Matsuoka, T., et al. (2015). Dysbiosis in the gut microbiota of patients with multiple sclerosis, with a striking depletion of species belonging to clostridia XIVa and IV clusters. PLoS One 10, 1–16. doi: 10.1371/journal.pone.0137429.

Mortzfeld, B. M., Taubenheim, J., Fraune, S., Klimovich, A. V, and Bosch, T. C. G. (2018). Stem Cell Transcription Factor FoxO Controls Microbiome Resilience in Hydra. 9, 1–10. doi: 10.3389/fmicb.2018.00629.

Müller, H. W. and R. F. and L. H. and K. (2022). Package “dplyr” Type Package Title A Grammar of Data Manipulation.

Naqib, A., Poggi, S., Wang, W., Hyde, M., Kunstman, K., and Green, S. J. (2018). Making and Sequencing Heavily Multiplexed,. Gene Expr. Anal. Methods Protoc. 1783, 149–169.

Needham, D. M., and Fuhrman, J. A. (2016). Pronounced daily succession of phytoplankton, archaea and bacteria following a spring bloom. Nat. Microbiol. 1, 16005. doi: 10.1038/nmicrobiol.2016.5.

Oksanen, J., Simpson, G. L., and Blanchet, F. G. (2022). Vegan: Community Ecology Package. R package version 2.5–7. Available at: https://cran.r-project.org/web/packages/vegan/vegan.pdf (Accessed April 17, 2022).

Ondov, B. D., Bergman, N. H., and Phillippy, A. M. (2011). Interactive metagenomic visualization in a Web browser. BMC Bioinformatics 12, 385. doi: 10.1186/1471-2105-12-385.

Parada, A. E., and Fuhrman, J. A. (2017). Marine archaeal dynamics and interactions with the microbial community over 5 years from surface to seafloor. ISME J. 11, 2510–2525. doi: 10.1038/ismej.2017.104.

Parada, A. E., Needham, D. M., and Fuhrman, J. A. (2016). Every base matters: Assessing small subunit rRNA primers for marine microbiomes with mock communities, time series and global field samples. Environ. Microbiol. 18, 1403–1414. doi: 10.1111/1462-2920.13023.

Peixoto, R. S., Rosado, P. M., Leite, D. C. de A., Rosado, A. S., and Bourne, D. G. (2017). Beneficial microorganisms for corals (BMC): Proposed mechanisms for coral health and resilience. Front. Microbiol. 8, 1–16. doi: 10.3389/fmicb.2017.00341.

Pollock, J., Glendinning, L., Wisedchanwet, T., and Watson, M. (2018). The madness of microbiome: Attempting to find consensus “best practice” for 16S microbiome studies. Appl. Environ. Microbiol. 84. doi: 10.1128/AEM.02627-17.

Quast, C., Pruesse, E., Yilmaz, P., Gerken, J., Schweer, T., Yarza, P., et al. (2013). The SILVA ribosomal RNA gene database project: Improved data processing and web-based tools. Nucleic Acids Res. 41, 590–596. doi: 10.1093/nar/gks1219.

Rosenberg, E., and Zilber-Rosenberg, I. (2011). Symbiosis and development: The hologenome concept. Birth Defects Res. Part C - Embryo Today Rev. 93, 56–66. doi: 10.1002/bdrc.20196.

Rosenberg, E., and Zilber-Rosenberg, I. (2018). The hologenome concept of evolution after 10 years. Microbiome 6, 78. doi: 10.1186/s40168-018-0457-9.

Salje, J. (2021). Cells within cells: Rickettsiales and the obligate intracellular bacterial lifestyle. Nat. Rev. Microbiol. 19, 375–390. doi: 10.1038/s41579-020-00507-2.

Schnedler-Meyer, N. A., Kiørboe, T., and Mariani, P. (2018). Boom and Bust: Life History, Environmental Noise, and the (un)Predictability of Jellyfish Blooms. Front. Mar. Sci. 5. doi: 10.3389/fmars.2018.00257.

Schön, M. E., Martijn, J., Vosseberg, J., Köstlbacher, S., and Ettema, T. J. G. (2022). The evolutionary origin of host association in an ancient bacterial clade. Nat. Microbiol., 1189–1199. doi: 10.1038/s41564-022-01169-x.

Stabili, L., Parisi, M. G., Parrinello, D., and Cammarata, M. (2018a). Cnidarian interaction with microbial communities: From aid to animal’s health to rejection responses. Mar. Drugs 16, 1–15. doi: 10.3390/md16090296.

Stabili, L., Parisi, M. G., Parrinello, D., and Cammarata, M. (2018b). Cnidarian interaction with microbial communities: From aid to animal’s health to rejection responses. Mar. Drugs 16. doi: 10.3390/md16090296.

Stabili, L., Rizzo, L., Fanizzi, F. P., Angilè, F., Coco, L. Del, Girelli, C. R., et al. (2019). The jellyfish rhizostoma pulmo (Cnidaria): Biochemical composition of ovaries and antibacterial lysozyme-like activity of the oocyte lysate. Mar. Drugs 17. doi: 10.3390/md17010017.

Takagi, T., Naito, Y., Inoue, R., Kashiwagi, S., Uchiyama, K., Mizushima, K., et al. (2019). Differences in gut microbiota associated with age, sex, and stool consistency in healthy Japanese subjects. J. Gastroenterol. 54, 53–63. doi: 10.1007/s00535-018-1488-5.

Thompson, A. L., Monteagudo-Mera, A., Cadenas, M. B., Lampl, M. L., and Azcarate-Peril, M. A. (2015). Milk- and solid-feeding practices and daycare attendance are associated with differences in bacterial diversity, predominant communities, and metabolic and immune function of the infant gut microbiome. Front. Cell. Infect. Microbiol. 5, 1–15. doi: 10.3389/fcimb.2015.00003.

Thompson, L. R., Sanders, J. G., McDonald, D., Amir, A., Ladau, J., Locey, K. J., et al. (2017). A communal catalogue reveals Earth’s multiscale microbial diversity. Nature 551, 457–463. doi: 10.1038/nature24621.

Tinta, T., Kogovšek, T., Klun, K., Malej, A., Herndl, G. J., and Turk, V. (2019a). Jellyfish-associated microbiome in the marine environment: Exploring its biotechnological potential. Mar. Drugs 17. doi: 10.3390/md17020094.

Tinta, T., Kogovšek, T., Klun, K., Malej, A., Herndl, G. J., and Turk, V. (2019b). Jellyfish-Associated Microbiome in the Marine Environment: Exploring Its Biotechnological Potential. Mar. Drugs 17, 94. Available at: internal-pdf://118.247.10.80/Tinta 2019 Jellyfish microbiome review.pdf.

Vacca, M., Celano, G., Calabrese, F. M., Portincasa, P., Gobbetti, M., and De Angelis, M. (2020). The controversial role of human gut lachnospiraceae. Microorganisms 8, 1–25. doi: 10.3390/microorganisms8040573.

Van De Water, J. A. J. M., Allemand, D., and Ferrier-Pagès, C. (2018). Host-microbe interactions in octocoral holobionts - recent advances and perspectives. Microbiome 6, 1–28. doi: 10.1186/s40168-018-0431-6.

van Oppen, M. J. H., and Blackall, L. L. (2019). Coral microbiome dynamics, functions and design in a changing world. Nat. Rev. Microbiol. 17, 557–567. doi: 10.1038/s41579-019-0223-4.

Vargas-Albores, F., Ortiz-Suárez, L. E., Villalpando-Canchola, E., and Martínez-Porchas, M. (2017). Size-variable zone in V3 region of 16S rRNA. RNA Biol. 14, 1514–1521. doi: 10.1080/15476286.2017.1317912.

Viver, T., Orellana, L. H., Hatt, J. K., Urdiain, M., Díaz, S., Richter, M., et al. (2017). The low diverse gastric microbiome of the jellyfish Cotylorhiza tuberculata is dominated by four novel taxa. Environ. Microbiol. Available at: internal-pdf://78.43.122.251/Viver 2017microbiomeCotylorhiza.pdf.

Voolstra, C. R., Suggett, D. J., Peixoto, R. S., Parkinson, J. E., Quigley, K. M., Silveira, C. B., et al. (2021). Extending the natural adaptive capacity of coral holobionts. Nat. Rev. Earth Environ. 2, 747–762. doi: 10.1038/s43017-021-00214-3.

Wear, E. K., Wilbanks, E. G., Nelson, C. E., and Carlson, C. A. (2018). Primer selection impacts specific population abundances but not community dynamics in a monthly time-series 16S rRNA gene amplicon analysis of coastal marine bacterioplankton. Environ. Microbiol. 20, 2709–2726. doi: 10.1111/1462-2920.14091.

Weiland-Bräuer, N., Pinnow, N., Langfeldt, D., Roik, A., Güllert, S., Chibani, C. M., et al. (2020). The Native Microbiome is Crucial for Offspring Generation and Fitness of Aurelia aurita. MBio 11, 1–20. doi: 10.1128/mBio.02336-20.

Wickham, H. (2016). ggplot2: Elegant Graphics for Data Analysis. Springer-Verlag New York. Available at: http://link.springer.com/10.1007/978-0-387-98141-3.

Willis, C., Desai, D., and Laroche, J. (2019). Influence of 16S rRNA variable region on perceived diversity of marine microbial communities of the Northern North Atlantic. FEMS Microbiol. Lett. 366, 1–9. doi: 10.1093/femsle/fnz152.

Yang, B., Wang, Y., and Qian, P. Y. (2016). Sensitivity and correlation of hypervariable regions in 16S rRNA genes in phylogenetic analysis. BMC Bioinformatics 17, 1–8. doi: 10.1186/s12859-016-0992-y.

Yeh, Y. C., McNichol, J., Needham, D. M., Fichot, E. B., Berdjeb, L., and Fuhrman, J. A. (2021). Comprehensive single-PCR 16S and 18S rRNA community analysis validated with mock communities, and estimation of sequencing bias against 18S. Environ. Microbiol. 23, 3240–3250. doi: 10.1111/1462-2920.15553.

Yilmaz, P., Parfrey, L. W., Yarza, P., Gerken, J., Pruesse, E., Quast, C., et al. (2014). The SILVA and “all-species Living Tree Project (LTP)” taxonomic frameworks. Nucleic Acids Res. 42, 643–648. doi: 10.1093/nar/gkt1209.

Zhang, S. L., Bai, L., Goel, N., Bailey, A., Jang, C. J., Bushman, F. D., et al. (2017). Human and rat gut microbiome composition is maintained following sleep restriction. Proc. Natl. Acad. Sci. U. S. A. 114, E1564–E1571. doi: 10.1073/pnas.1620673114.

Zilber-Rosenberg, I., and Rosenberg, E. (2008). Role of microorganisms in the evolution of animals and plants: The hologenome theory of evolution. FEMS Microbiol. Rev. 32, 723–735. doi: 10.1111/j.1574-6976.2008.00123.x.

