## Supplementary figures and images for "Selecting 16S rRNA primers for microbiome analysis in a host-microbe system: the case of the jellyfish *Rhopilema nomadica*"

### Supplementary Figure 2

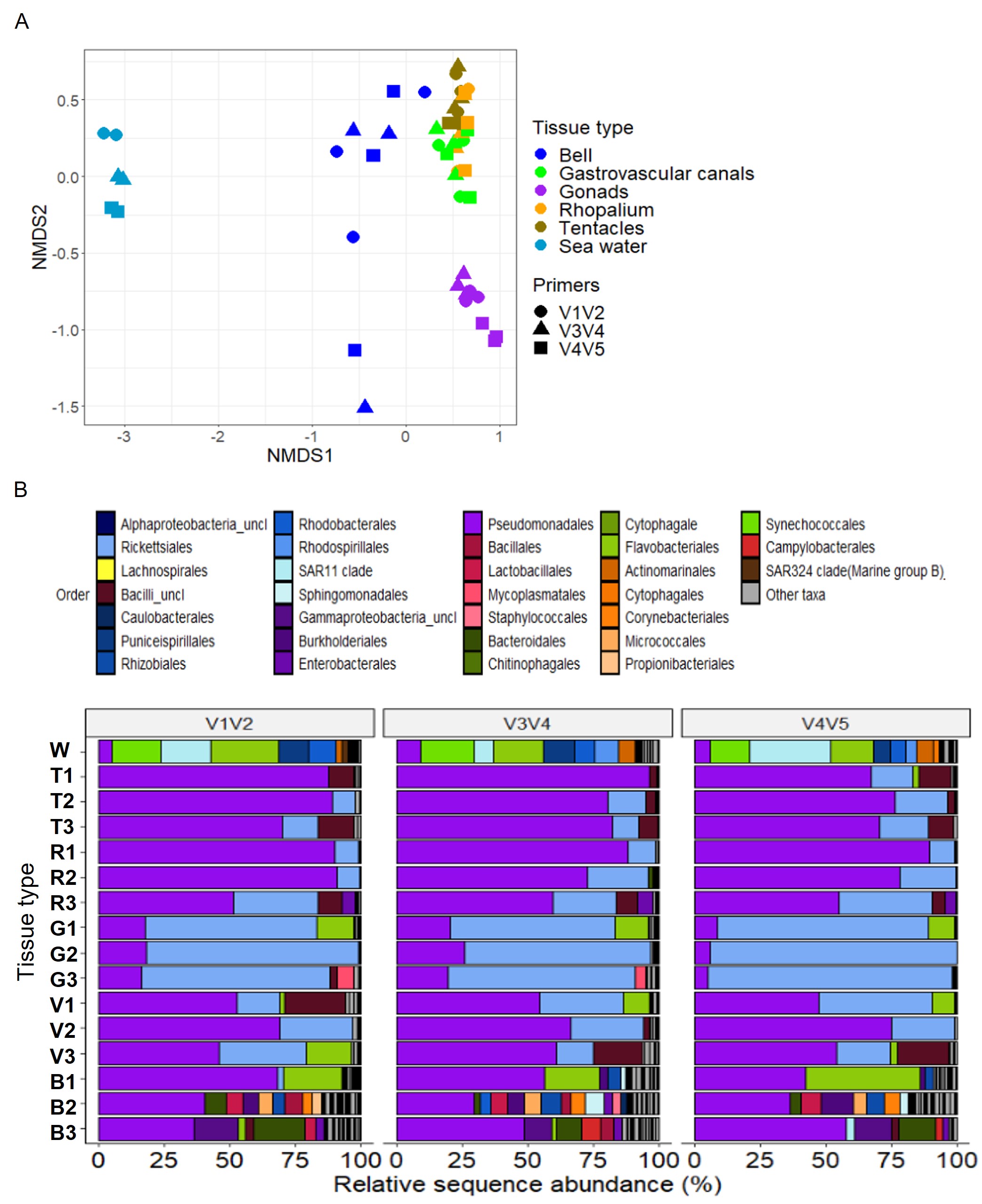

### Supplementary Figure S1

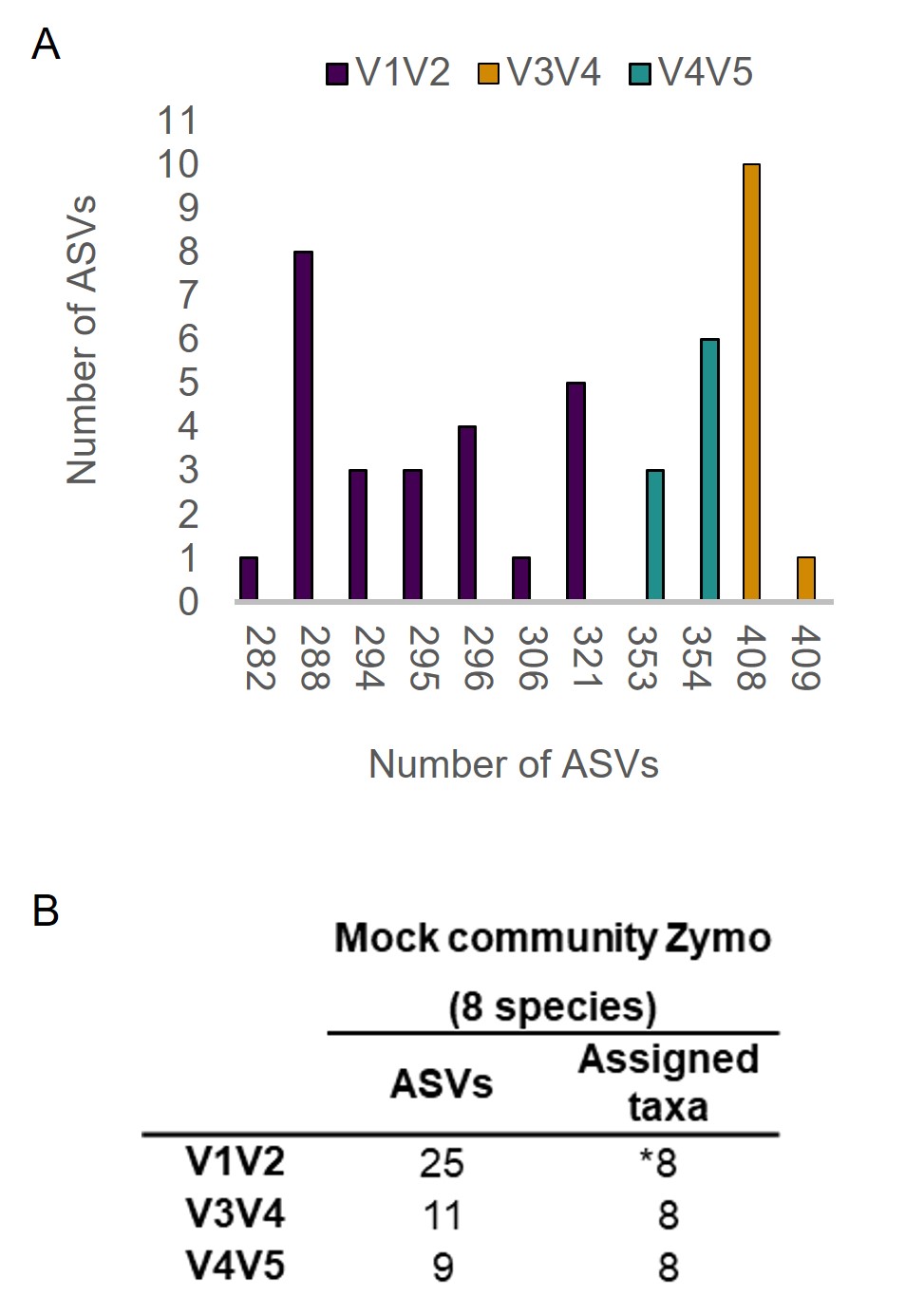
